# Regulation of an antibiotic resistance efflux pump by quorum sensing and a TetR-family repressor in *Chromobacterium subtsugae*

**DOI:** 10.1101/2023.09.02.556004

**Authors:** Pratik Koirala, Cassie Doody, Helen Blackwell, Josephine R. Chandler

## Abstract

The soil bacterium *Chromobacterium substugae* uses a single LuxI-R-type quorum-sensing system, CviI-R, to regulate genes in a cell density-dependent manner. CviI synthesizes the signal *N*-hexanoyl-homoserine lactone (C6-HSL) and CviR is a C6-HSL-responsive cytoplasmic transcription regulator. C6-HSL-bound CviR activates dozens of genes, for example the *cdeAB-oprM* cluster coding for an efflux pump conferring antibiotic resistance. The *cdeAB-oprM* genes are also regulated by an antibiotic-responsive transcription factor, CdeR, which represses expression of these genes. We are interested in understanding how *C. subtsugae* integrates different environmental cues to regulate antibiotic resistance. In this study, we sought to delineate the mechanism of regulation of the *cdeAB-oprM* genes by CviR and CdeR. In recombinant *E. coli,* the *cdeA* promoter is activated by CviR and repressed by CdeR. We identify non-overlapping sequence elements in the *cdeA* promoter that are required for CviR activation and CdeR repression, respectively. We also examined the role of CdeR in modulating *cdeA* activation by C6-HSL in *C. subtsugae*. We show that CviR and CdeR can independently modulate transcription from the *cdeA* promoter in *C. subtsugae*, consistent with the conclusion that CviR and CdeR regulate the *cdeAB-oprM* genes by interacting directly with different binding sites in the *cdeA* promoter. These results contribute to a molecular understanding of how the *cdeAB-oprM* genes are regulated and provide new insight into how *C. subtsugae* integrates different environmental cues to regulate antibiotic resistance.

**Importance:** Many bacteria regulate antibiotic resistance in response to antibiotics and other cues from the environment. In many cases the regulatory mechanisms are best understood in the context of clinical isolates where mutations frequently emerge in resistance regulation pathways. However, an understanding of the role of antibiotic resistance regulators in integrating environmental information is less well understood. In the soil bacterium *Chromobacterium subtsugae,* an antibiotic-resistance gene cluster is regulated by population density and antibiotics through two different transcription factors; the quorum sensing signal receptor CviR and an antibiotic-responsive transcription factor CdeR. In this study, we show that these factors independently modulate the transcription of the antibiotic resistance genes and coordinate to ensure sensitive responses to changes in cell density. The results give new insight into antibiotic resistance regulation in *C. subtsugae* and contribute to a broader understanding of how bacteria optimize the regulation of antibiotic resistance in response to changes in the environment.

## Introduction

Many proteobacteria use quorum sensing to sense changes in population density and coordinate population-wide changes in gene regulation. One type of quorum sensing in proteobacteria involves N-acyl-homoserine lactone (AHL) signals. AHLs are produced by LuxI-family synthases. They are secreted at low levels and diffuse in and out of the cells so that their environmental concentration increases with increasing population density [1]. When the AHLs reach a critical threshold in the environment they bind to LuxR-family signal receptors, which are cytoplasmic transcription regulators [2]. AHL binding causes conformational changes in the LuxR protein that enables them to activate the transcription of target genes [3]. LuxR-dependent gene activation typically requires binding at a conserved 16- to 20-bp DNA sequence in the promoter of the target genes, which is called a *lux* box [4].

Many genes that are activated by quorum sensing are also regulated by other transcription factors. For example, in some cases gene activation requires quorum sensing and also another transcription regulator, similar to an AND logic gate where two inputs are needed to turn on a singular output [5]. In this way the bacterial population can integrate information from population density and other environmental inputs to modulate changes in gene expression. For example, hydrogen cyanide production in *Pseudomonas aeruginosa* is activated by quorum sensing and the anaerobic regulator (Anr) [6]. In this way the bacterial population can modulate expression of hydrogen cyanide in response to several environmental inputs. Such regulation could be important for adapting quickly to changing conditions.

Our group has focused on the nonpathogenic saprophyte *Chromobacterium subtsugae* as a model for studying quorum sensing because it has a single quorum-sensing circuit and has standard growth kinetics and nutritional requirements that are useful for interspecies interaction studies. The *C. subtsugae* quorum-sensing system is CviR-I [7]. CviI synthesizes *N-*hexanoyl homoserine lactone (C6-HSL), and C6-HSL-bound CviR binds to a canonical *lux-*box-like sequence in the promoter of target genes [8]. Activation of the CviR-I system induces expression of dozens of genes [9]. Many of the CviR-I-responsive genes have roles that are predicted or known to be important for competing with other bacteria, for example by activating the production of the purple-pigmented antibiotic violacein [7] and hydrogen cyanide [9].

Our prior work showed that *C. subtsugae* also uses CviR-I to increase resistance to several antibiotics by transcriptionally activating genes coding for a resistance-nodulation division (RND)-family efflux pump, CdeAB-OprM [10, 11]. RND pumps use energy to transport substrates, such as antibiotics, out of the cytoplasm [12]. These pumps have a periplasmic membrane fusion protein (CdeA), which forms a tripartite complex with a periplasmic adaptor protein (CdeB), and an outer membrane protein (OprM). Upstream of the *cdeAB-oprM* genes and facing the opposite direction is a gene coding for a predicted TetR-family repressor called CdeR. In previous studies we confirmed that CdeR represses *cdeAB-oprM* expression [11], although it was unknown whether this regulation was direct or indirect through another regulatory pathway. In addition the potential interaction of CdeR with the quorum sensing activator CviR remained unknown.

In this study, we sought to elucidate a molecular understanding of *cdeAB-oprM* gene regulation by the two regulators, CviR and CdeR. We used heterologous *E. coli* to demonstrate that each of these regulators can modulate the expression of the promoter controlling the *cdeAB-oprM* genes (P*cdeA*). We mutationally map the required DNA elements in P*cdeA* for regulation by each regulator and validate the CviR binding site using purified protein *in vitro.* Our findings demonstrate that the DNA elements for CviR and CdeR are non-overlapping, suggesting these two regulators can have independent effects on regulating transcription from P*cdeA*. We confirm the independent regulatory role of each of these regulators by monitor the expression of a *PcdeA-lacZ* transcriptional reporter in *C. subtsugae* and use this reporter to confirm that the independent effects of CviR and CdeR. We also show that the presence of CdeR maximizes the range of signal responsiveness and sensitivity of CviR. Our results add new insight into the function of the CdeR and CviR regulators and establish a system for future studies to understand how these regulators optimize antibiotic resistance in *C. subtsugae*.

## Materials and methods

### Bacterial culture conditions

All strains were grown in Luria-Bertani (LB) broth, in LB containing morpholinepropanesulfonic acid (LB-MOPS; 50 mM; pH7), or on LB with 1.5%(wt/vol) Bacto agar. All broth cultures were incubated with shaking at 30°C (*Chromobacterium subtsugae*) or 37°C (*Escherichia coli*). Synthetic acyl-homoserine lactone signals were purchased from Cayman Chemicals (MI, USA) and stored in ethyl acetate acidified with 0.01% of glacial acetic acid. In all cases, synthetic signals were added to the culture flask and dried down completely using a stream of nitrogen gas before adding the culture media. When required, antibiotics were added in the following concentrations: carbenicillin 100 µg ml^-1^, tetracycline 40 µg ml^-1^, gentamicin 50 µg ml^-1^ for *C. subtsugae* and carbenicillin 100 µg ml^-1^, tetracycline 20 µg ml^-1^, gentamicin 20 µg ml^-1^ for *E. coli*.

Genomic DNA, PCR and DNA fragments, and plasmid DNA were purified according to the manufacturer’s protocol by using a Puregene Core A kit, plasmid purification miniprep kit, or PCR cleanup/gel extraction kits (Qiagen or IBI-MidSci) respectively. RNA was isolated using the Qiagen RNEasy Minikit. β-galactosidase activity was measured using the Tropix Galacto-Light Plus™ chemiluminescent kit according to the manufacturer’s protocol (Applied Biosystems, Foster City, CA).

### Bacterial strains and plasmids

Bacterial strains, plasmids, and oligonucleotides used in this study are summarized in Table 1. *C. subtsugae* Cv017 (referred to as wild type, previously known as *C. violaceum* CV017) is the derivative of strain ATCC 31532 with a transposon insertion in gene CV_RS05185 causing activation of violacein production [7]. For plasmid construction, we used *E. coli* strain DH10B (Invitrogen).

**Table 1.**
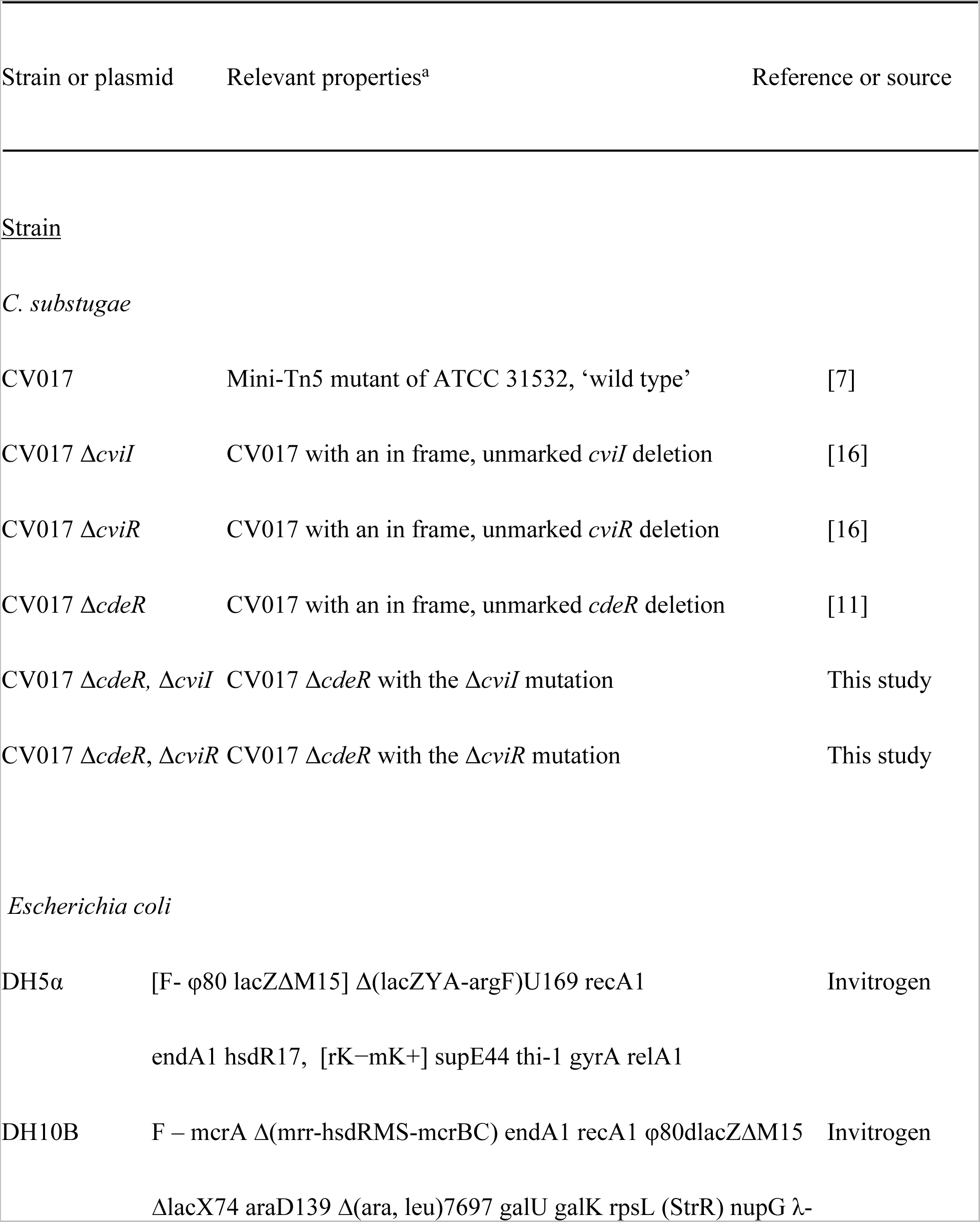
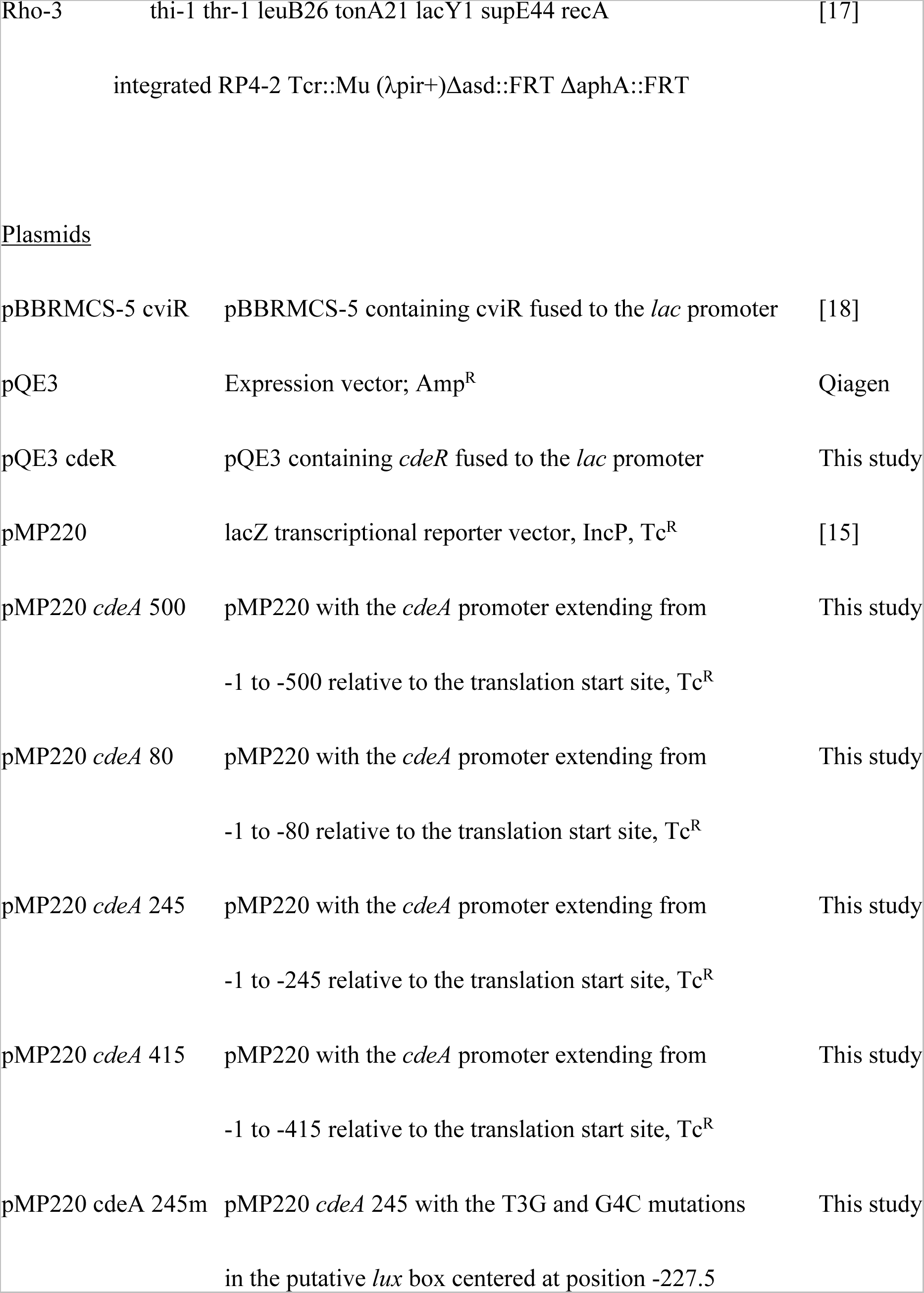
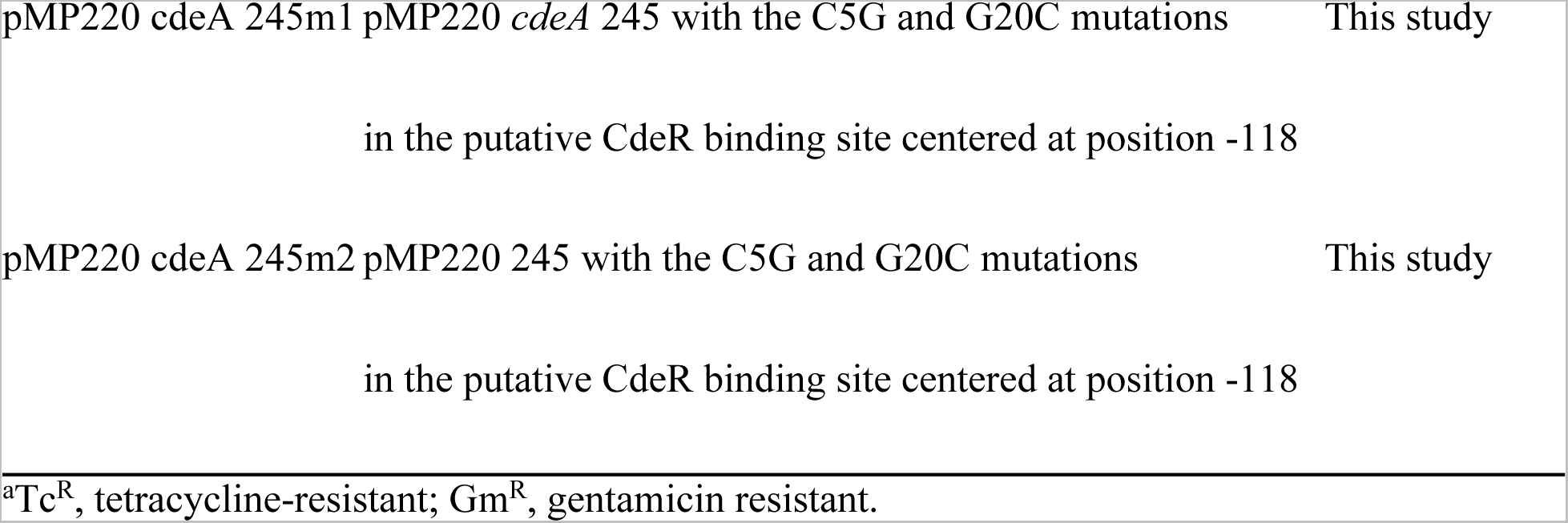
Bacterial strains and plasmids used in this study.

*C. subtsugae* mutants were constructed using allelic exchange as described previously [13]. Briefly, gene deletions were introduced to *C. subtsugae* by conjugation using the suicide delivery plasmid pEXG2 [14]. Merodiploids were selected on LB agar containing gentamicin, and deletion mutants were counter-selected on LB + 15% sucrose. Mutant strains were verified by testing for gentamicin sensitivity and by PCR amplification and sequencing of the deletion region.

For transcription reporter assays, plasmid pMP220 with a DNA fragment spanning 500 bp upstream of the translation start site of *cdeA* (pMP220 *cdeA-lacZ*) or with the mutated or truncated *cdeA* promoter regions were made by either PCR amplification or by DNA synthesis (Genscript, USA). The DNA products were made with incorporated *kpnI* or *xbaI* enzyme recognition sites and were digested with KpnI and XbaI and ligated to KpnI-XbaI-digested pMP220 [15] upstream of the *lacZ* reporter. The *cdeR* expression plasmid pQE30 *cdeR* was made by amplifying the *cdeR* gene with primer-incorporated BamHI and HindIII restriction enzyme sites, digesting with BamHI/HindIII and ligating to BamHI/HindIII-digested pQE30 (Qiagen).

### Transcription reporter assays

To assess activation of the *cdeA* promoter in recombinant *E. coli,* we used *E. coli* strain DH5a carrying *cviR* downstream of the *lac* promoter in pBBRMCS-5 (pBBRMCS-5 *cviR*) and pMP220 carrying the full-length, truncated or mutated *cdeA* promoter fused to the *lacZ* reporter gene. In some cases, we added a third plasmid, pQE30 *cdeR.* In all cases, overnight cultures were used as starters by diluting them to an OD_600_ of 0.05. After 3 h shaking at 30°C, the cultures were added to 2 ml Eppendorf tubes containing dried AHLs as indicated. The volume of culture in each tube was 0.5 ml. After 6 more h shaking at 30C, β-galactosidase activity was measured as described above.

To assess activation of the *cdeA* promoter in *C. subtsugae, C. subtsugae* strains carrying the pMP220 *cdeA* 450 reporter plasmid were used as starters by diluting to an OD_600_ of 0.05. When experimental cultures reached an OD_600_ of 0.5 they were distributed to 2 ml Eppendorf tubes containing different concentrations of dried AHLs. The volume in each tube was 0.5 ml. After 5 h with shaking at 30°C, b-galactosidase activity was measured using the Tropix Galacto-Light Plus™ chemiluminescent kit according to the manufacturer’s protocol (Applied Biosystems, Foster City, CA).

### Droplet digital PCR

RNA was harvested from *C. subtsugae* grown in LB-MOPS to an OD of 2. RNA was prepared using the RNeasy minikit (Qiagen) by following the manufacturer’s instructions. Droplet digital PCR (ddPCR) was performed on a Bio-Rad QX200 ddPCR system using Eva Green supermix. Each reaction mixture contained 1 ng μl−1 of cDNA template, 0.25 μM each primer, 10 μl Eva Green supermix, and 8 μl H2O in a 20-μl volume. After generating 40 μl of oil droplets, 40 rounds of PCR were conducted using the following cycling conditions: 94°C for 20 s, 60°C for 20 s, and 72°C for 20 s. Absolute transcript levels were determined using Bio-Rad QuantaSoft software. In all cases, a no-template control was run with no detectable transcripts. The reference gene used was glyceraldehyde-3-phosphate dehydrogenase (GAPDH, encoded by the *gapA* gene), and the results are reported as the calculated transcript amount of a given gene per calculated *gapA* transcript.

## Results

### The *cdeA* promoter is activated by CviR and C6-HSL in heterologous *E. coli*

To assess activation of the *cdeA* promoter by CviR in *E. coli,* we engineered a plasmid with a 500-bp fragment containing the promoterless *lacZ* fused to the presumed *cdeA* promoter (from positions −1 to −500 with respect to the *cdeA* translation start codon) and introduced this plasmid to heterologous *Escherichia coli* carrying a second plasmid with *cviR* expressed from a constitutively activated *lac* promoter. In *E. coli,* activation of the *cdeA::lacZ* reporter was dependent on CviR and C6-HSL (Fig. 1A). The concentration of C6-HSL required for half-maximal *cdeA* promoter activation was 19 nM, which is similar to the responses observed for other LuxR homologs to their cognate AHLs [19–21]. This result supports the view that CviR directly activates the *cdeA* promoter.

**Fig. 1.**
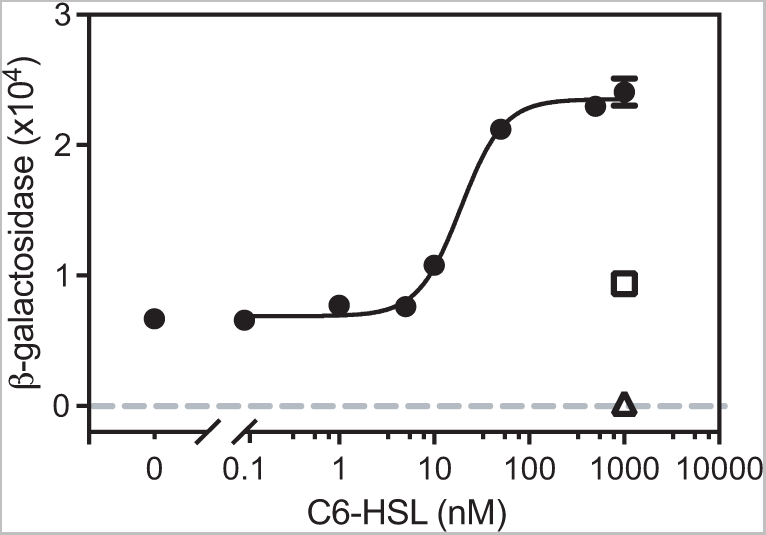
Expression of *cdeA* requires CviR and C6-HSL in recombinant *E. coli*. *E. coli* carried a CviR expression vector (pBBRMCS-5 *cviR*) and a *cdeA-lacZ* reporter (pMP220 *cdeA* 500) (filled circles connected by a fitted curve). ≤, control showing the response of *E. coli* with the reporter plasmid and pBBRMCS-5 vector with no CviR. Δ, control showing the response of *E. coli* with the CviR expression vector and the pMP220 vector with the promoterless *lacZ.* Values are the mean of three independent experiments and the vertical lines indicate standard deviations. The fitted curve was generated using a nonlinear regression model with variable slope using Prism 9.5.0 (R^2^ = 0.9929 and degrees of freedom = 20).

### CviR activation of the *cdeA* promoter requires a *lux* box-like sequence

In the DNA region upstream of the *cdeA* promoter, we identified three potential *lux* boxes with sequences resembling the canonical *lux* box sequences identified in *Chromobacterium violaceum,* a related species [8](Fig. 2A). These putative *lux* box-like sequences are centered at −72.5, −227.5 and −253.5 relative to the translation start site (Fig. 2A). To determine if any of these sequences are important for CviR to activate *cdeA* transcription, we engineered additional *cdeA* promoter-reporter fusions starting from different sites in the 5’ end of the promoter. The longest *cdeA* promoter fragment started at position −277 relative to the *cdeA* translation start site and contained all three of the identified *lux* box-like sequences. With this fragment, C6-HSL increased transcription of the *lacZ* reporter ∼3.5-fold (Fig. 2B), similar to that of the original 500 bp fragment a smaller *cdeA* promoter fragment starting at position −245 that did not contain the −253.5 *lux* box-like sequence showed similar C6-HSL-dependent activation as that of the −500 and −277 fragments, supporting that the −253.5 *lux-*box-like fragment is not required for CviR activation of *cdeA*. However, a shorter fragment starting at base pair −219 that was missing the −227.5 *lux-*box-like fragment was not activated by C6-HSL (Fig. 2B). These results upport the conclusion that the *lux* box-like sequence centered at position −227.5 is required for C6-HSL-dependent activation of the *cdeA* promoter by CviR. Interestingly, the 219-bp fragment had almost no measurable *lacZ* activity in the absence of C6-HSL, compared with the low-level basal activation observed with the longer fragments (Fig. 2B). These results support the idea that the transcription start site might be encoded within the region between −219 and −245 relative to the *cdeA* translation start site. The placement of the transcription start site in this region is consistent with a *lux* box-like element centered at position −227.5.

**Fig. 2.**
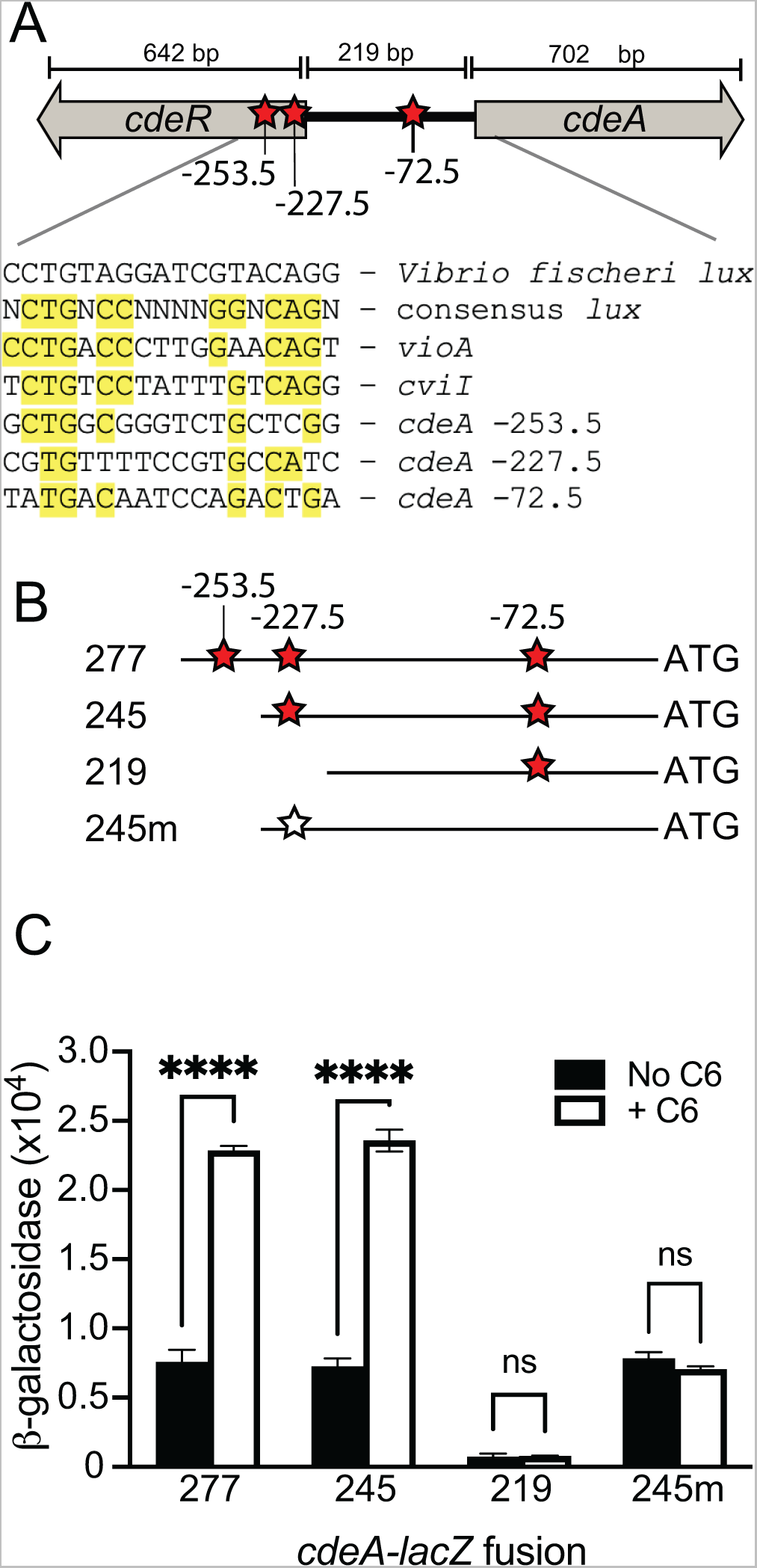
Identification of the site required for CviR to activate transcription from the *cdeA* promoter. A. Map showing the divergently transcribed *cdeR-cdeA* region. Red stars indicate putative *lux* boxes, with *lux* box-like sequenced indicated by the position in the DNA relative to the *cdeA* translation start site. The sequence of each putative *lux* box aligned with the *V. fischeri lux* box, the *C. violaceum* consensus *lux* box and the experimentally identified *lux* box within the promoter of the *C. violaceum vioA* and *C. subtsugae cviI* is shown below the map. B. Schematic of *cdeA-lacZ* promoter regions used for analysis in part C. The *cdeA* translation start site is shown on the far right (3’ end, ‘ATG’). The numbers indicate the base position with respect to the *cdeA* translation start site where the 5’ end of the fragment starts. The fragment labeled “m” has mutations in the *lux* box-like element (T3C and G4A) as indicated by the open star. C. Transcription from different fragments containing the *cdeA* promoter was measured in recombinant *E. coli* carrying the pMP220 plasmid with *cdeA* fragments as indicated and the CviR expression plasmid pBBRMCS-5 CviR. 1 μM C6-HSL was added as indicated. Statistical significance by two-way ANOVA using Sidak’s multiple comparisons test; **** p<0.001 (ns, not significant).

To test the hypothesis that the *lux* box-like element centered at position −227.5 is needed for CviR to activate the *cdeA* promoter, we mutated the T and G residues at the 3rd and 4th position of this sequence within the *cdeA* fragment starting at position −245 (T3G and G4C, named “-245m” in Fig. 3B). Introducing these mutations to the *lux* box-like element abolished C6-HSL-dependent activation of the *lacZ* reporter and had no effect on reporter activation in the absence of C6-HSL (Fig. 2C). These results suggest the mutations disrupted the ability of CviR to activate this promoter without disrupting promoter recognition by the RNA polymerase.

**Fig. 3.**
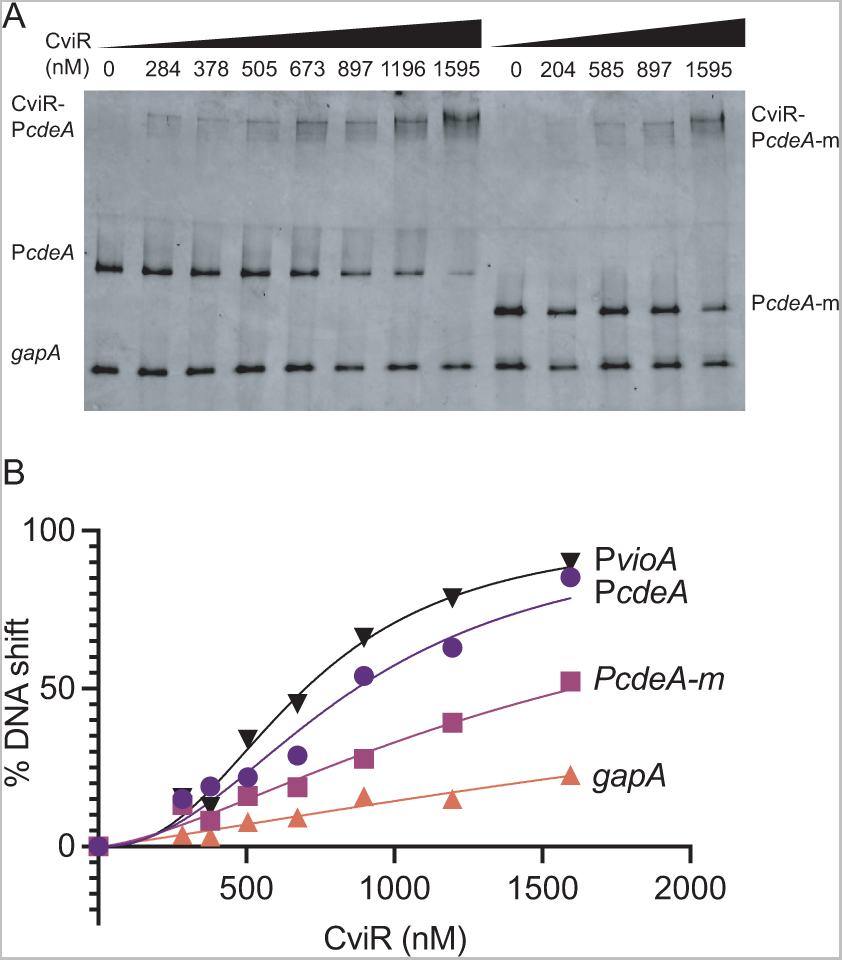
Binding of purified CviR to *cdeA* promoter DNA containing the *lux* box-like element. (A) Representative DNA mobility shift assay. Lane 1 to 8 contained WT *cdeA* target DNA. Lane 9-13 contained a mutated *cdeA* target DNA with TG to GA substitution in the *lux-box*. All lanes also contained a non-specific probe (*gapA* coding sequence). The molar amount of CviR in each binding reaction is indicated. All binding reactions contained 5 μM C6-HSL. B) Graphical representation of % DNA shift from Figure 4A. Data are the average of three independent experiments.

To study the interaction of CviR with the *cdeA* promoter, we purified CviR as an N-terminal histidine fusion protein from recombinant *E. coli* and used the purified protein to perform gel shift experiments (Fig. 3). When incubated with 5 μM C6-HSL, CviR bound to the 245-bp promoter DNA fragment extending from positions −245 to −1 relative to the *cdeA* translation start site. Binding was dose-dependent. This fragment contains the *lux* box-like element centered at position −227.5 and identical to the −245 *cdeA* promoter fragment indicated in Fig. 3B. At concentrations above These results support the conclusion that CviR interacts directly with the *cdeA* promoter and requires the *lux* box-like element at −227.5 for this interaction.

### CdeR represses transcription from the *cdeA* promoter in heterologous *E. coli*

Although *cdeA* transcription is repressed by CdeR, we created a plasmid encoding *cdeR* fused downstream of the IPTG-inducible *lac* promoter. We introduced this plasmid or the pQE3 vector control to heterologous *E. coli* carrying the 500-bp *cdeA-lacZ* reporter and CviR expression plasmids. We compared *cdeA-lacZ* activation in the presence or absence of CdeR and with 100 or 500 nM C6-HSL or no signal (**Fig. 5A**). In the absence of signal we saw no effects of CdeR. However, in the presence of either signal concentration, CdeR significantly decreased *cdeA-lacZ* activity compared with cells with no CdeR. This result supports the idea that CdeR directly represses the activation of the *cdeA* promoter, possibly by blocking transcription.

The DNA recognition sites of TetR family proteins can be quite divergent. Thus, we used the protein fold recognition tool Phyre2 [22] to identify proteins with high structural relatedness to CdeR (Table S1). The top two hits were *Pseudomonas aeruginosa* TtgR [23] and *Escherichia coli* AcrR [24]. These two proteins have similar DNA recognition sequences (Fig. 5A). We searched the DNA sequence upstream of the *cdeA* translation start site for a region with similarity to the TtgR and AcrR recognition motifs and found a sequence with a high degree of similarity to the TtgR sequence. This sequence was centered at base −118 relative to the CdeA translation start site (Fig. 4A and B). To test the importance of this sequence for CdeR to repress *cdeA* activation, we generated *cdeA* promoter fragments with two or three point mutations in conserved residues previously shown to be important for TtgR binding [24]. One fragment (m1) had mutations C5G and G20C, and a second fragment (m2) had the C5G and G20C mutations and an additional mutation T23A. These mutated fragments were fused to *lacZ* in plasmid pMP220 and the plasmids were introduced to our *E. coli* strain carrying the CviR and CdeR expression plasmids. Results with these plasmids showed that in the presence of 1 𝜇M C6-HSL, CdeR poorly suppressed *lacZ* activation in the m1 fragment and no longer repressed *lacZ* activation in the m2 fragment (Fig. 4C). These results support the conclusion that the recognition site located at −118 relative to the *cdeA* translation start site is required for CdeR to repress activation of the *cdeA* promoter.

**Fig. 4.**
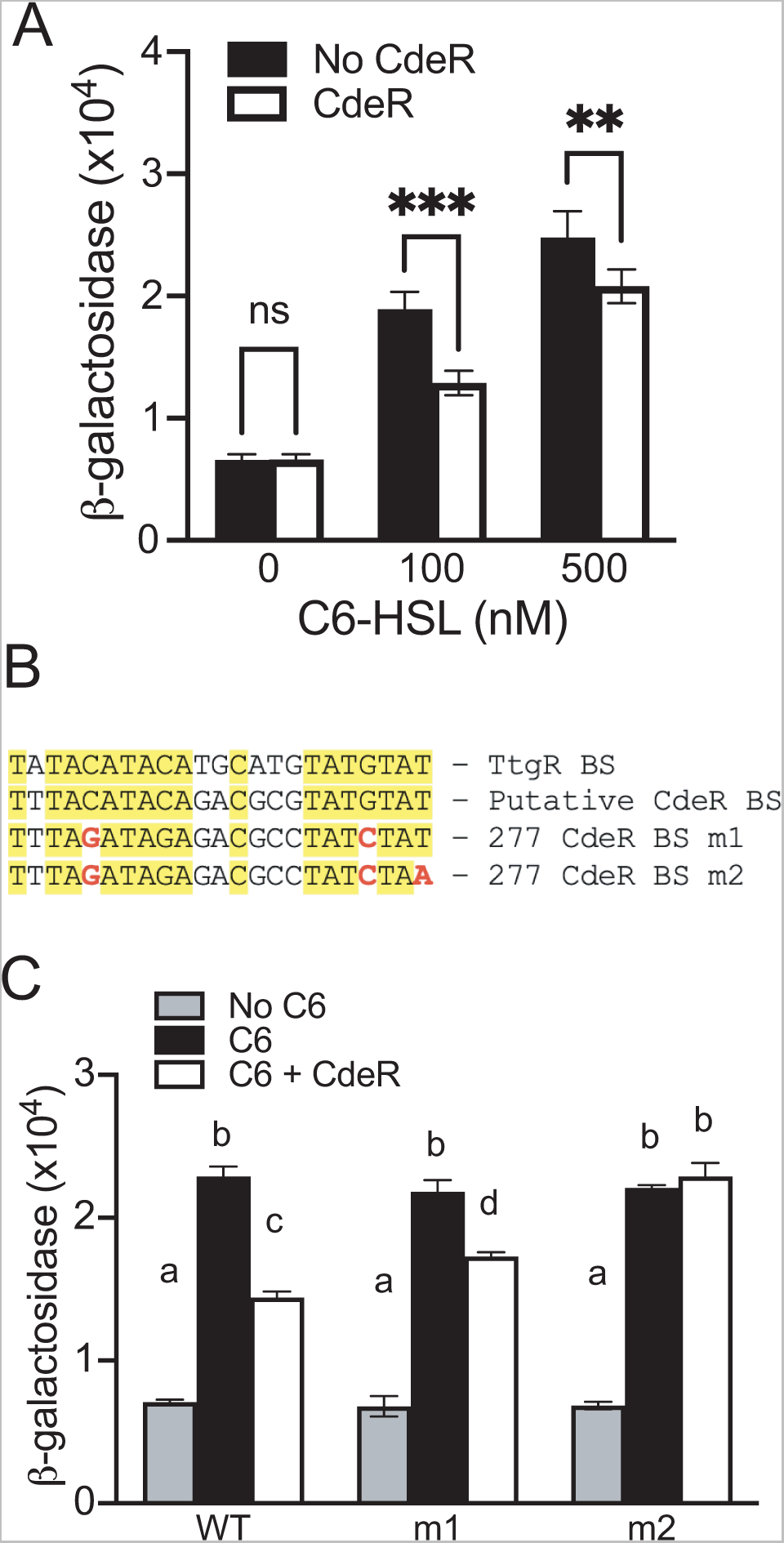
CdeR repression of *cdeA* promoter activation. A. Transcription from the *cdeA* promoter was measured in recombinant *E. coli* carrying the P*cdeA-lacZ* reporter plasmid pMP220 *cdeA* 500, pBBRMCS-5 CviR, and pQE3 *cdeR* or the empty pQE3 vector. 1 μM C6-HSL was added as indicated. Statistical significance by two-way ANOVA using Sidak’s multiple comparisons test; **** p<0.001. B. The sequence of the *Pseudomonas putida* TtgR binding site (BS) aligned with the putative CdeR binding site and the two mutated CdeR binding sites used in this study (m1 and m2). C. Transcription activation of *cdeA-lacZ* as in part A. WT, P*cdeA* sequence unchanged; m1, P*cdeA* carrying the C5G and G20C mutations in the putative CdeR binding site; m2, P*cdeA* carrying the mutations in m1 and an additional T23A mutation.. Statistical significance by two-way ANOVA using Tukey’s multiple comparisons test with different letters indicating p < 0.05 and same letters indicating p > 0.05.

### Regulation of *cdeA* transcription by CviR and CdeR in *C. subtsugae*

Our results support a model where CviR and CdeR interact with nonoverlapping elements in the *cdeA* promoter to regulate expression of this gene (Figs. 5A and S1). The CviR binding site overlaps with the translation start site of CdeR, which is upstream from *cdeA* and facing in the opposite direction. Although CviR binding in this region could possibly block CdeR transcription, we did not observe any effects of CviR on *cdeR* transcription in *C. subtsugae* Fig. S2).

Based on our model, we predicted that CdeR and CviR could modulate transcription of *cdeA* independently in *C. subtsugae*. To test this prediction, we compared *cdeA* transcripts in wild-type (CV017) and strains carrying single deletions of *cviR* or *cdeR* as well as the doubleΔ*cdeR-cviR* mutant. We reasoned that if CdeR and CviR are functionally independent, they should each be able to regulate *cdeA* in the presence or absence of the other regulator. In support of this model, deleting *cdeR* increased the level of *cdeA* transcripts in the presence or absence of *cviR* (Fig. 5B, compare WT withΔ*cdeR* andΔ*cviR* withΔ*cviR-cdeR*). In addition, deleting *cviR* decreased the level of *cdeA* transcripts in the presence or absence of *cdeR* (Fig. 5B, compare WT with Δ*cviR* and Δ*cdeR* with Δ*cviR-cdeR*). These results support the conclusion that CviR and CdeR can each regulate *cdeA* independently in *C. subtsugae*.

**Fig. 5.**
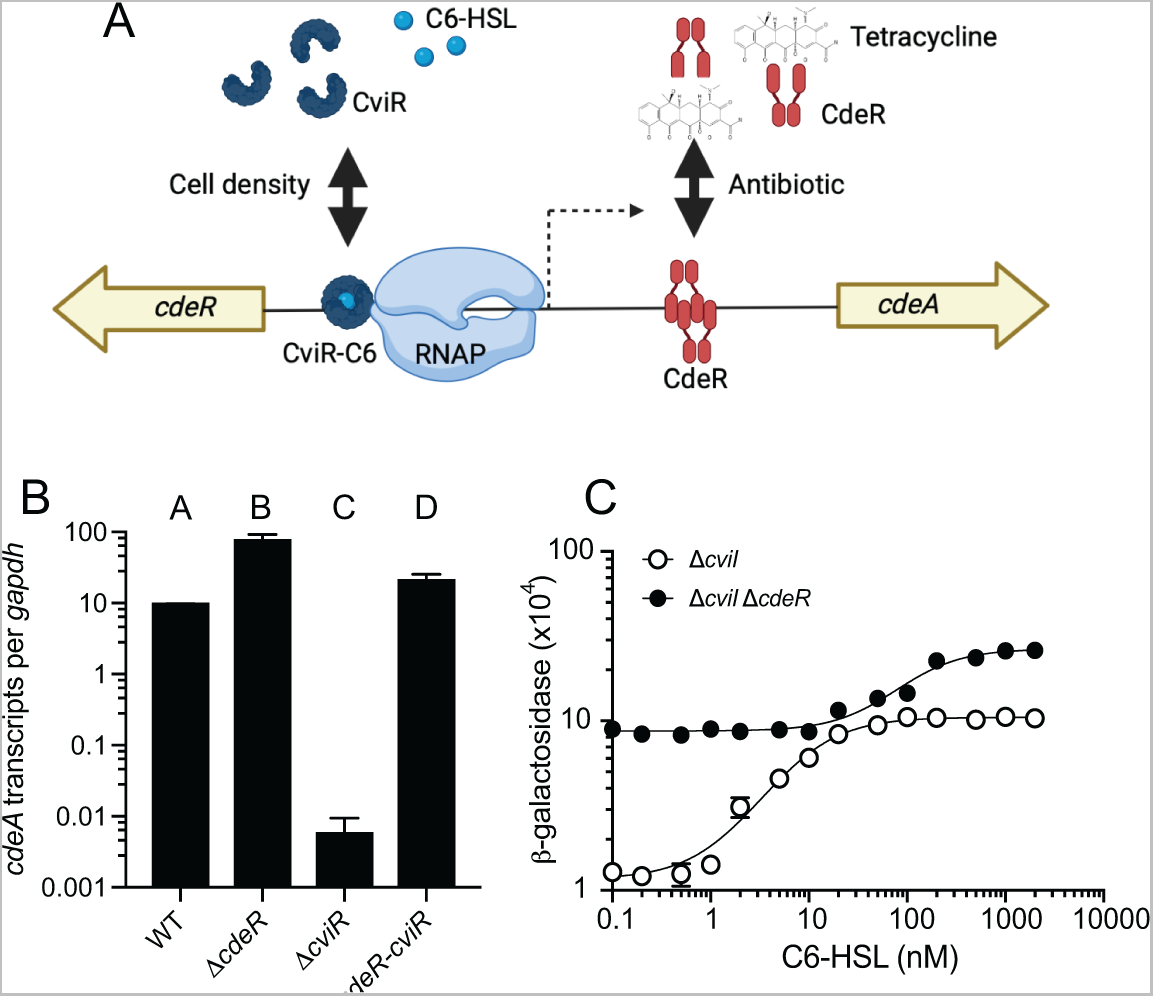
Regulation of *cdeA* and *cdeR* transcription by CviR and CdeR in *C. subtsugae*. A. Model of CviR and CdeR regulation of the *cdeA* promoter. CviR is a C6-HSL-responsive activator and CdeR is an antibiotic (tetracycline)-responsive repressor. B. Transcripts were measured from exponentially growing cells using quantitative reverse-transcription PCR (qRT-PCR) using primers specific to the open reading frame of *cdeA* (A and C) or *cdeR* (B). Results are shown as *gapdh-*adjusted transcript levels. The values represent the means of three independent experiments and the vertical bars represent the standard deviation of the means. Statistical analysis by one-way ANOVA showed significant differences (p>0.01) in comparing A-B, C-D, A-C and B-D (see text). C. Activation of the *cdeA::lacZ* reporter plasmid in the *C. subtsugae ΔcviI* single mutant or the Δ*cviI-cdeR* double mutant in response to increasing concentrations of C6-HSL. Results are the average of 3 independent experiments; error bars show the standard deviation. In most cases the error is not larger than the symbol and is not visible.

Although CviR and CdeR function independent, we observed that the magnitude of regulation varied depending on the presence or absence of the other regulator. This result suggested there may be some interactive effects between these two regulators. To address this possibility, we assessed whether the presence or absence of the CdeR repressor altered the *cdeA* promoter sensitivity or magnitude change in response to AHL activation. We moved our *cdeA::lacZ* reporter fusion plasmid into the *C. subtsugae* Δ*cviI* single mutant or the Δ*cviI-cdeR* double mutant and compared reporter activation in response to increasing concentrations of C6-HSL. In both strains, we observed a C6-HSL-dependent increase in *cdeA-lacZ* reporter activity, consistent with our studies of *cdeA* gene transcription (Fig. 5B). However, the concentration of C6-HSL required for half-maximal *cdeA* promoter activation was higher in the absence of CdeR compared with the presence of CdeR. In the presence of CdeR we also observed a greater magnitude change in response to C6-HSL compared with no CdeR (9-fold vs. 3-fold). These results suggest the presence of the CdeR repressor modulates CviR sensitivity and magnitude of activation by C6-HSL.

## Discussion

The results of this study provide information to contribute to our molecular understanding of how *C. subtsugae* regulates the *cdeAB-oprM* genes. Regulation involves two transcription factors, the quorum sensing activator CviR and the antibiotic-responsive repressor CdeR. The results support the idea that the *cdeA* promoter is directly regulated by each of these regulators through binding at non-overlapping regulatory sequences in the promoter DNA. We also show that there are interactive effects of CdeR on the ability of CviR to respond to signals. These results provide new insight into how *C. subtsugae* regulates gene transcription in response to population density and antibiotics. The results also contribute to a broader understanding of how bacteria can coordinately control gene expression in response to changing environments.

Our results show that CviR interacts with a *lux* box-like element upstream of the predicted transcription start site in the *cdeA* promoter (Fig. 5A). This site is positioned at –227.5 relative to the CdeA translation start site. The CviR-binding *lux* box-like element likely overlaps with the −35 recognition sequence of RNAP, similar to other LuxR-family proteins [25]. CdeR likely recognizes a sequence downstream from this site, centered at base −118. The positioning of the *lux* box-like element and −35 RNAP recognition sequence suggest the transcription start site is likely positioned around −200, which places this sequence upstream from the CdeR recognition sequence. This suggests CdeR might repress activation from the *cdeA* promoter by blocking transcription elongation or “roadblocking” [26, 27]. Such regulation has been previously described for other proteins, such as the catabolite control protein CcpA in *Bacillus subtilis* [28], the purine biosynthesis regulator PurB in *Escherichia coli* [29], and by synthetically re-positioning the binding site of other DNA repressors, such as the *E. coli lac* repressor LacI (e.g. [30–32]) . In some cases, roadblocking can be overcoming by increasing the level of transcription initiation [26]. In this manner, CviR might help to overcome a CdeR-dependent roadblock by increasing the activation of transcription from the *cdeA* promoter.

The *cdeAB-oprM* genes encode an efflux pump that can increase antibiotic resistance by pumping antibiotics out of the cell. What could be the evolutionary advantage of regulating the expression of CdeAB-OprM by both CdeR and CviR? Regulation that enables the gene to respond to two different environmental inputs simultaneously might be important to fine-tune the levels of expression in response to different environments. Such regulation might help to balance the costs of efflux pump production with the benefits in certain environments. There might also be other benefits of coordinating expression by both regulators, for example by modulating the coordination of efflux pump production across a population of cells. For example, roadblocking by CdeR could possibly increase the stochasticity of efflux pump expression in the population, causing some cells to become highly antibiotic resistant while others remain susceptible. Stochastic gene expression can help a population diversify costly traits as a strategy of risk-spreading or bet hedging [33]. Regulation by multiple transcription factors could provide an opportunity to increase or decrease bet hedging strategies in response to different environments.

Our studies of *cdeA* regulation in the native *C. subtsugae* provide mechanistic insight into the function of CdeR and CviR. Our results showed that the presence of CdeR increased signal sensitivity and dynamic range of response by CviR. The change in signal sensitivity might be because CdeR increases basal levels of promoter activation, which masks low signal responses. In the case of the dynamic range of signal responses, a previous study examining the dynamic range of ligand inducible promoter activation might provide insight. This study demonstrated that changes in the strength of promoter binding to the RNA polymerase can modulate the dynamic range of ligand responsiveness [34]. CdeR likely reduces the apparent promoter strength by blocking transcription elongation, which in turn enhances the dynamic range. The role of promoter binding strength in modulating dynamic range of signal responses suggest that the *cdeA* promoter might have evolved to have an optimally tuned promoter such that CdeR binding enhances the dynamic range. Such binding might maximize the ability to sense and respond to population density in the absence of antibiotic.

## Acknowledgements

This work was supported by the National Institutes of Health (NIH) R35GM133572 to J.R.C. and R35GM131817 to H.B. and by the KU Weaver Fellowship to P.K.

