## Supplemental file for "Regulation of an antibiotic resistance efflux pump by quorum sensing and a TetR-family repressor in *Chromobacterium subtsugae*"

Table S1. Top hits for CdeR using Phyre2 structure-based alignment tool[a].

| Protein Name | Organism Name | %ID[b] | Alignment Coverage |
|---|---|---|---|
| TtgR | *Pseudomonas putida* | 27 | 95% from residue 2-206 |
| AcrR | *Escherichia coli K-12* | 28 | 95% from residue 6-210 |
| NalD | *Pseudomonas aeruginosa* PAO1 | 36 | 92% from residue 8-205 |
| YfiR | *Bacillus subtilis* | 12 | 95% from residue 2-206 |
| HapR | *Vibrio cholera* | 16 | 92% from residue 1-198 |

[a]Alignments were done using the the normal modeling mode function of Phyre2 (1).
[b]% ID refers to the percent amino acid identity overall.

**GCGAGGTGCGCGACACGCCCTGCTCGCTGAACAGCCGCTCGGCGGCATCCAGCAGCAGCTG**

CviR binding site

**GCGGGTCTGCTCGGCCTCTTCACGTGTTTTCCGTGCCAT**CCATCATTCTTCGCTCTTAGTA

← Start of CdeR

CTAGTATGAATTTTAACGGTTCTTTGTTGCCCAAACCGCTGCGTTTTTACATATCCGAGAC

CdeR binding site

ATTATAGCATTTTACATACAGACGCGTATGTATGTAAAATACCGAAGCTTCAGGTTATGAC

AATCCAGACTGACCATCCCCTGAAGCCCTATCCATTTGATTACCTTCCCGATTTTGCTTTC

Start of CdeA →

GAGGATTCGATACC**ATGCAAGGA**...

**Figure S1. Sequence of the promoter region of the *C. subtsugae* efflux pump gene *cdeA*** The sequence is from strain Cv017, a derivative of ATCC31532 (accession NZ_LKIW00000000). The coding region of CdeA is bolded in blue and that of CdeR is bolded in green (CdeR is in the reverse direction). The putative CviR and CdeR binding sites determined in this manuscript are highlighted by red boxes.

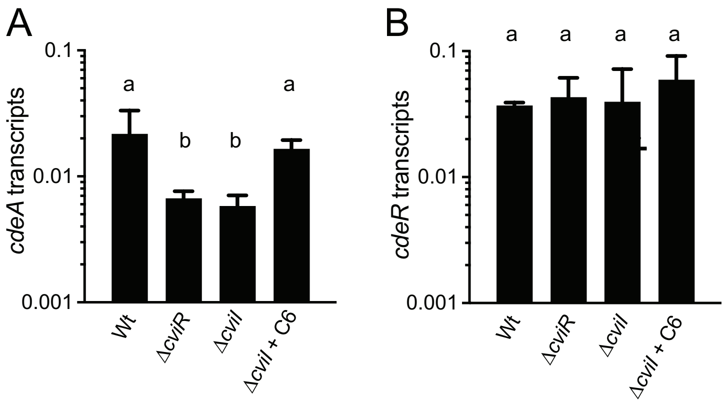


**Fig. S2. Regulation of *cdeA* and *cdeR* transcription by CviR and CdeR.** Transcripts were measured from exponentially growing cells using quantitative reverse-transcription PCR (qRT-PCR) using primers specific to the open reading frame of *cdeA* (A) or *cdeR* (B). Results are shown as *gapdh*-adjusted transcript levels. The values represent the means of three independent experiments and the vertical bars represent the standard deviation of the means. Statistical analysis by one-way ANOVA showed significant variation among means of the data shown in A ($p<0.02$) but not B ($p>0.5$).
